# An aneuploidy epistasis map reveals metabolic vulnerabilities associated with supernumerary chromosomes in cancer

**DOI:** 10.1101/2024.09.30.615609

**Authors:** R. Y. Magesh, A. N. Kaur, F. N. Keller, A. Frederick, T. Tseyang, J. A. Haley, A. M. Rivera-Nieves, A. C. Liang, D. A. Guertin, J. B. Spinelli, S. J. Elledge, E. V. Watson

## Abstract

Despite the general detriment of aneuploidy to cellular fitness, >90% of solid tumors carry an imbalanced karyotype. Regardless of this existing paradox, our understanding of the molecular responses to aneuploidy remains limited. Here, we explore these cellular stresses and unique vulnerabilities in aneuploid human mammary epithelial cells (HMECs) enriched for breast cancer-associated copy number alterations (CNAs). To uncover the genetic dependencies specific to aneuploid cells, we conducted a comprehensive, genome-wide CRISPR knockout screen targeting isogenic diploid and aneuploid HMEC lines. Our study reveals that aneuploid HMECs exhibit an increased reliance on pyrimidine biosynthesis and mitochondrial oxidative phosphorylation genes, and demonstrate heightened fitness advantages upon loss of tumor suppressor genes. Using an integrative multi-omic analysis, we confirm nucleotide pool insufficiency as a key contributor to widespread cellular dysfunction in aneuploid HMECs with net copy number gain. While diploid cells can switch seamlessly between pyrimidine synthesis and salvage, cells with increased chromosomal content exhibit p53 activation and S-phase arrest when relying on salvage alone, and exhibit increased sensitivity to DNA-damaging chemotherapeutics. This work advances our understanding of the consequences of aneuploidy and uncovers potential avenues for patient stratification and therapeutic intervention based on tumor ploidy.

## Introduction

Aneuploidy - the state of having abnormal numbers of chromosomes - is a hallmark of cancer, affecting >90% of tumors, with most tumor genomes falling in the >2N to <4N ploidy range^1^ and displaying tissue-specific karyotype selection patterns^2^. Aneuploidy has been proposed to provide selective proliferative or survival benefit during tumorigenesis^3,4^ by shaping tumor suppressor and oncogene dosage^5,6^, however aneuploidy also induces cellular stress and is generally detrimental to cellular fitness^3,7–11^. How tumor cells overcome the burdens associated with an imbalanced genome, often a net excess of chromosomes, is not well understood but may represent exploitable nodes of vulnerability in cancer^12–15^.

The burden of excess chromosomes in net-gain aneuploidy induces stress across a myriad of biochemical systems that synthesize the polymers of life: DNA, RNA, and protein. Just how widespread these stresses permeate across cellular systems is still being uncovered, with documentation of significant disruption to replication^16–18^, transcription^3,19–21^, RNA metabolism^22^, autophagy^23,24^ and proteostasis^8,25,26^. In yeast, drosophila, and mouse embryonic fibroblasts, chromosomal imbalance results in metabolic disruptions in the form of increased ROS production^27–29^. It is no surprise then that aneuploidy generally slows growth rates in experimental models ranging from yeast to human cell lines, with the exception of some recurrent copy number alterations found prevalently in cancer, which have been shown to increase growth rate in cognate tissue-of-origin models^3^.

Aneuploidy is associated with p53 activation^28,30^, which has been proposed to derive from aneuploidy-associated replication stress^17^. This can lead to growth arrest, senescence, or apoptosis^31–33^. Thus, one function of p53 is to sense and restrict aneuploidy in the body, and loss of p53 by somatic mutation in tumors is strongly associated with increased levels of aneuploidy compared to tumors without p53 mutations^34–36^. In fact, patients with Li Fraumeni syndrome who harbor germline p53 mutations have increased levels of aneuploidy in tissues and develop many different types of cancer - often in the same patient^37^. p53 is the top growth-restricting factor in aneuploid RPE1 cells, as determined through unbiased functional genomic methods^22,38^. However, the underlying mechanisms by which aneuploidy leads to replication stress and subsequent p53 activation are poorly understood.

Here we utilize a functional genomics approach to characterize gene essentiality in net-gain aneuploid human mammary epithelial cells compared to isogenic diploid controls. By performing genome-wide CRISPR screens^39,40^ in paired aneuploid/diploid mammary cell lines^3^ and integrating transcriptomic, metabolomic, and bioenergetics data, we identify nucleotide pool insufficiency as a key underlying mechanism of p53 activation in net copy number gain aneuploid cells. Additionally, aneuploid cells exhibited increased mitochondrial ATP production, as well as increased glycolytic rates, to support the increased metabolic demands associated with increased chromosomal content. Nucleotide pool insufficiency associated with net-gain chromosomal aneuploidy could potentially be targeted therapeutically to enhance current chemotherapeutic modalities of breast cancer treatment, or stratify patients under current clinical standards of treatment.

## Results

### Genome-wide aneuploidy synthetic lethality CRISPR screens

With the goal of fully profiling aneuploidy synthetic lethalities in a cancer-relevant system, we performed genome-wide CRISPR screens in isogenic diploid and aneuploid cell lines derived from diploid hTERT-immortalized human mammary epithelial cells (hTERT-HMECs, referred to as “HMECs” here), which are considered good models for the cell type of origin of multiple breast cancer subtypes^3,41,42^. Additionally, HMECs are untransformed and do not contain driver mutations as assessed by deep whole genome sequencing^3^, enabling us to isolate the aneuploidy synthetic lethality profile without the complications of background mutational status. We chose four different aneuploid clonal lines with largely non-overlapping chromosomal amplifications, except for chromosome 20 gain which is recurrently selected in HMECs *in vitro*^3^ and is present in each line. We performed control screens utilizing three different clonally-derived diploid lines, also in duplicate, for a total of 14 genome-wide CRISPR screens (**Fig. 1 a**). This library contains 5 guides per gene targeting ∼18,000 protein-coding genes, for a total of ∼90,000 reagents.

**FIGURE 1:**
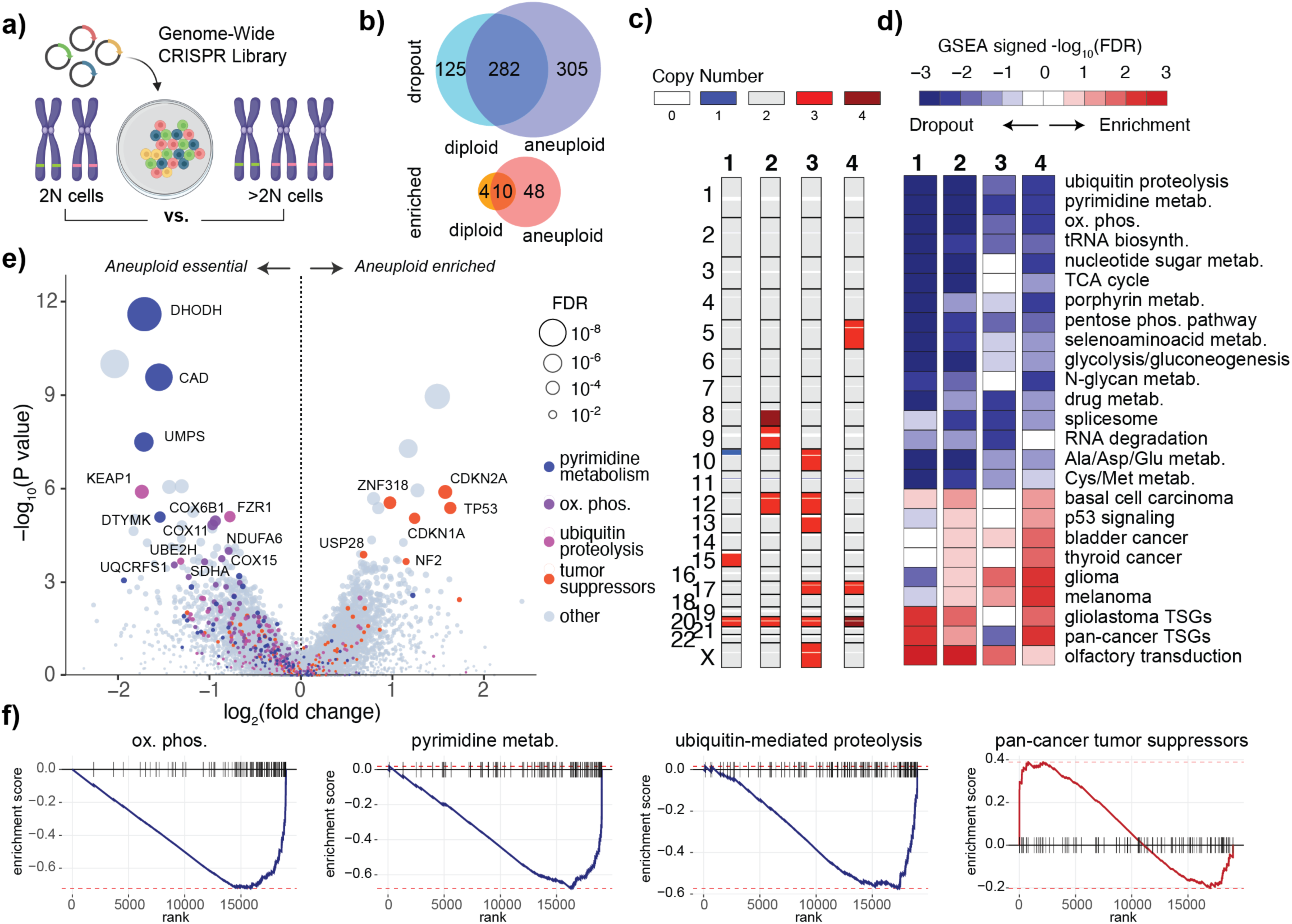
Net-gain aneuploidy epistasis profile in human mammary epithelial cells. **a,** Diagram of concept for genome-wide aneuploidy CRISPR screens. Four unique aneuploid HMEC clones with mostly non-overlapping CNAs were used for the screen alongside a paired, isogenic diploid control. **b,** Diagram showing the shared and unique differentially enriched/depleted genes between isogenic aneuploid and diploid HMECS **c,** Bar columns indicate the copy number profiles of each of the four unique aneuploid clones. **d,** Top enriched/depleted pathways across each of the four aneuploid clone screens, each compared to their isogenic diploid control **e,** Volcano plot showing gene-level meta-analysis of all aneuploid vs diploid screens, with specific genes colored based on their pathway involvement in pyrimidine metabolism, oxidative phosphorylation, ubiquitin-mediated proteolysis. Enriched tumor-suppressor genes are also labeled. **f,** Enrichment plots for oxidative phosphorylation genes, pyrimidine biosynthesis genes, ubiquitin-mediated proteolysis genes, and pan-cancer tumor suppressor genes in aneuploid HMEC screens compared to diploid control screens.

After infection with the Lentiviral CRISPR library at a multiplicity of infection of 0.3 and representation of 500 cells per guide, the initial “population doubling 0” (PD0) timepoint was collected, then cells were passaged for 6 population doublings and collected for the endpoint of the screen (PD6). Gene dropout/enrichment was determined based on sequencing coverage of guides after amplicon-seq of extracted genomic DNA using edgeR^43,44^. A slight bias in dropout/enrichment based on chromosomal location for genes on chromosomes affected by aneuploidy was observed as previously described^45^, which we implemented a computational correction to resolve (**Fig. S1 a**).

A start-to-endpoint analysis of aneuploid and diploid screens revealed largely overlapping gene dropout and enrichment profiles (**Fig. 1 b** and **Fig. S1 b**), however more genes were essential in aneuploid HMECs compared to diploid HMECs (FDR < 0.05 and log_2_ fold change < -2). To generate aneuploidy epistasis profiles, we compared PD6 endpoints between the aneuploid and diploid screens (**Fig. 1 c-f**). Gene set enrichment analysis of epistasis profiles for each of the four aneuploid mutants used in the screens revealed differential dropout of guides targeting genes in pathways known to be stressed in aneuploidy, like ubiquitin mediate proteolysis (**Fig. 1 d-f**). Additionally, we observe significantly stronger enrichment of p53 pathway-targeting guides (as well as other tumor suppressor pathways) in aneuploid lines compared to diploids (**Fig. 1 d-f, and Fig. S1 b, c**). Top-25 enriched hits in 3 out of 4 screens included prominent tumor suppressors TP53, CDKN1A, USP28, NF1 and CDKN2A (**Fig. S1 c**).

### Net-gain aneuploid cells do not tolerate *de novo* pyrimidine synthesis disruption and are insufficiently rescued with uridine

Top differential dropout hits in the aneuploidy screens centered around nucleotide biosynthesis and mitochondrial oxidative phosphorylation metabolic pathways (**Fig. 1 d-f**). The three enzymes that catalyze the first six reactions of the *do novo* pyrimidine biosynthesis pathway (CAD, DHODH, UMPS), were among the top 4 dropout hits (**Fig. 1 e and Fig. S1 d**). DHODH is positioned in the mitochondrial membrane and catalyzes the vital pyrimidine biosynthesis reaction of dihydroorotate (DHO) reduction to orotate, passing reducing equivalents directly to the mitochondrial ubiquinone pool^46,47^ **(Fig. 2 a)**. Thus, nucleotide metabolism and mitochondrial redox biology are inextricably linked, and this critical node renders mitochondrial oxidative phosphorylation indispensable for tumor cell proliferation despite high flux and ATP production through glycolysis^48,49^. Our data suggests this pyrimidine biosynthesis/mitochondrial oxidative phosphorylation axis is especially critical for proliferation of cells with supernumerary chromosomal content, perhaps due to increased nucleotide and energy requirements.

**FIGURE 2:**
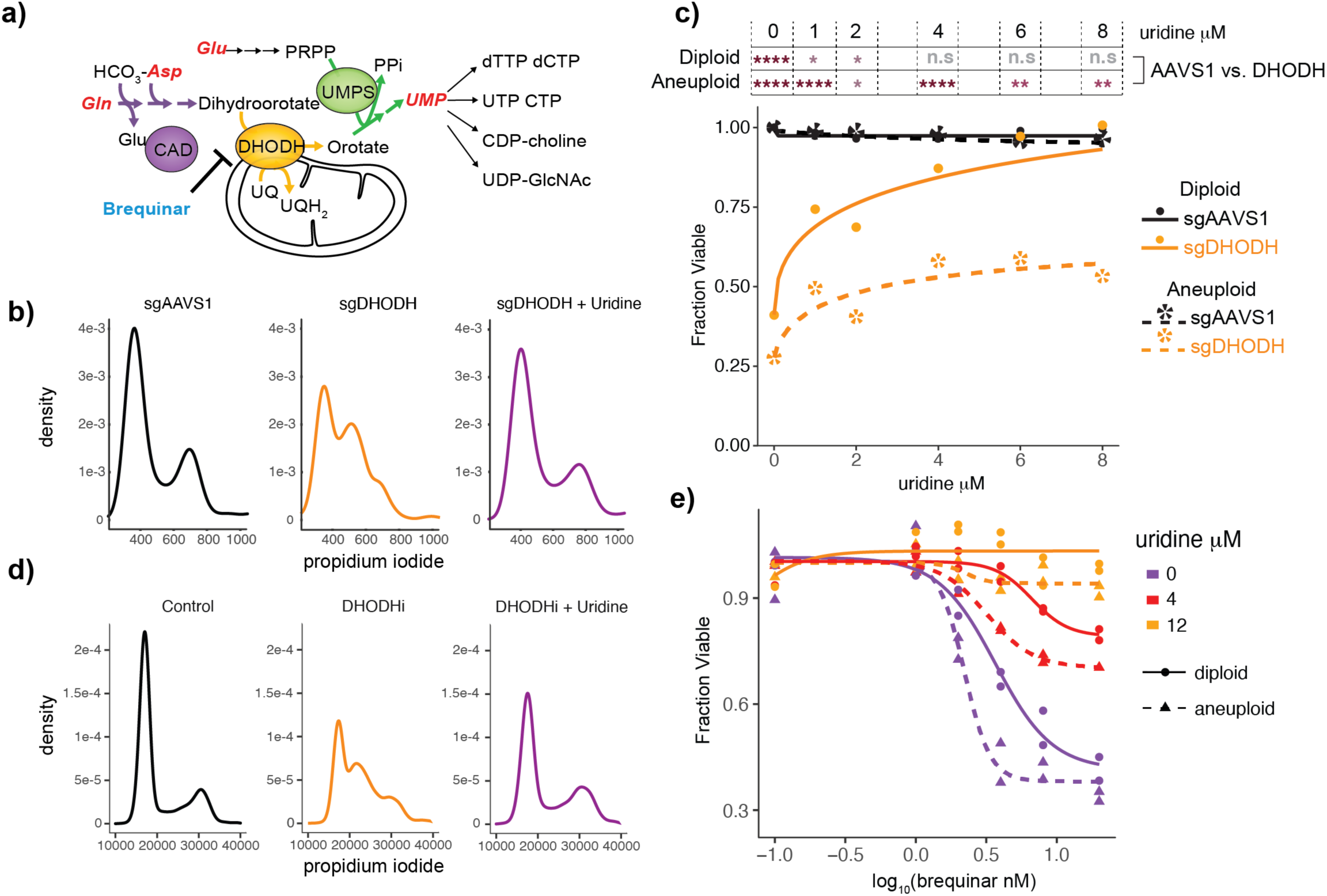
Net-gain aneuploid cells are sensitive to pyrimidine synthesis disruption. **a,** Diagram of pyrimidine biosynthesis pathway. **b,** Density plots showing DNA stained with propidium iodide in parental HMECs with AAVS1 (control) knockout, DHODH knockout, and DHODH knockout supplemented with uridine. DHODH knockout causes cells to arrest in S-phase, which is rescued by uridine. **c,** Cell viability of AAVS1 and DHODH knockout HMECs in diploid and net-gain aneuploid backgrounds under uridine supplementation ranging from 0-8 μM. The significance of the difference between the AAVS1 knockout and DHODH knockout cell viability in aneuploid and diploid backgrounds was calculated with T tests at each uridine dose assayed and is indicated by stars at the top of the figure. **** P-value < 0.0001; *** P-value < 0.001; ** P-value < 0.01; * P-value < 0.05. The aneuploid HMEC line used for this experiment is clone 2 in Fig. 1c. **d,** The DHODH inhibitor (DHODHi) brequinar phenocopies DHODH knockout. **e,** Cell viability of diploid (solid line/circles) and aneuploid (dotted line/ triangles) HMECs after treatment with brequinar doses ranging from 0.1-40 nM and uridine doses at 0, 4, and 12 μM (depicted using either purple, red, and yellow respectively).

To validate this dependency, we utilized CRISPR to mutate DHODH in representative aneuploid and diploid mammary epithelial cell lines. During the process of selection and recovery we supplemented cells with uridine to prevent cell cycle arrest/death. Complete protein loss in each line was confirmed via western blot (**Fig. S2 a**). DHODH knockout resulted in S-phase accumulation when uridine was withdrawn from culture in both diploid and aneuploid lines (**Fig. 2 b-c**). While the cell cycle arrest caused by DHODH mutation in diploid cells could be fully rescued by supplementing physiological plasma uridine concentrations (3-8 μM^50^), DHODH knockout in aneuploid cells could not be fully rescued by the same dose range of uridine supplementation (**Fig. 2 c**, **Fig. S2 b**). This indicates that net-gain aneuploid cells do indeed require more pyrimidine than diploid cells to replicate a higher chromosome load. This also indicates that, while diploid mammary epithelial cells are capable of fully sustaining growth using either nucleotide synthesis or nucleotide salvage alone, net-gain aneuploid cells cannot proliferate at full capacity using only pyrimidine salvage. This phenomenon could also be recapitulated with the DHODH inhibitor (DHODHi) brequinar^51^, which only mildly impaired diploid cell growth but severely impaired aneuploid cell growth under the same uridine supplementation conditions (**Fig. 2 d, e and Fig. S2 c**). Aneuploid cells entering S-phase arrest induced by DHODHi were less likely than diploid cells to successfully re-enter and progress through the cell cycle when provided uridine after arrest (**Fig. S2 d-f**).

### Net-gain aneuploid cells display insufficient pyrimidine synthesis relative to ploidy

To profile the metabolic constraints that may lead to nucleotide pool insufficiency in aneuploid HMECs, we traced de novo pyrimidine synthesis flux into UTP using ^13^C_4_-labeled aspartate revealed that net gain aneuploid HMECs do not increase pyrimidine synthesis capacity, displaying equal label incorporation into UTP compared to diploids (**Fig. 3 a, b**). However, since aneuploid HMECs have increased DNA content, this equivalent flux of de novo nucleotide synthesis compared to diploid cells represents a possible replication vulnerability, given that pyrimidine synthesis-to-ploidy ratios are effectively decreased (**Fig. 3 c-e**). Additionaly, we observe a significant decrease in GSH::GSSG ratios, indicating a disruption of redox balance and increased reactive oxygen species in aneuploid HMECs (**Fig. 3 f**).

**FIGURE 3:**
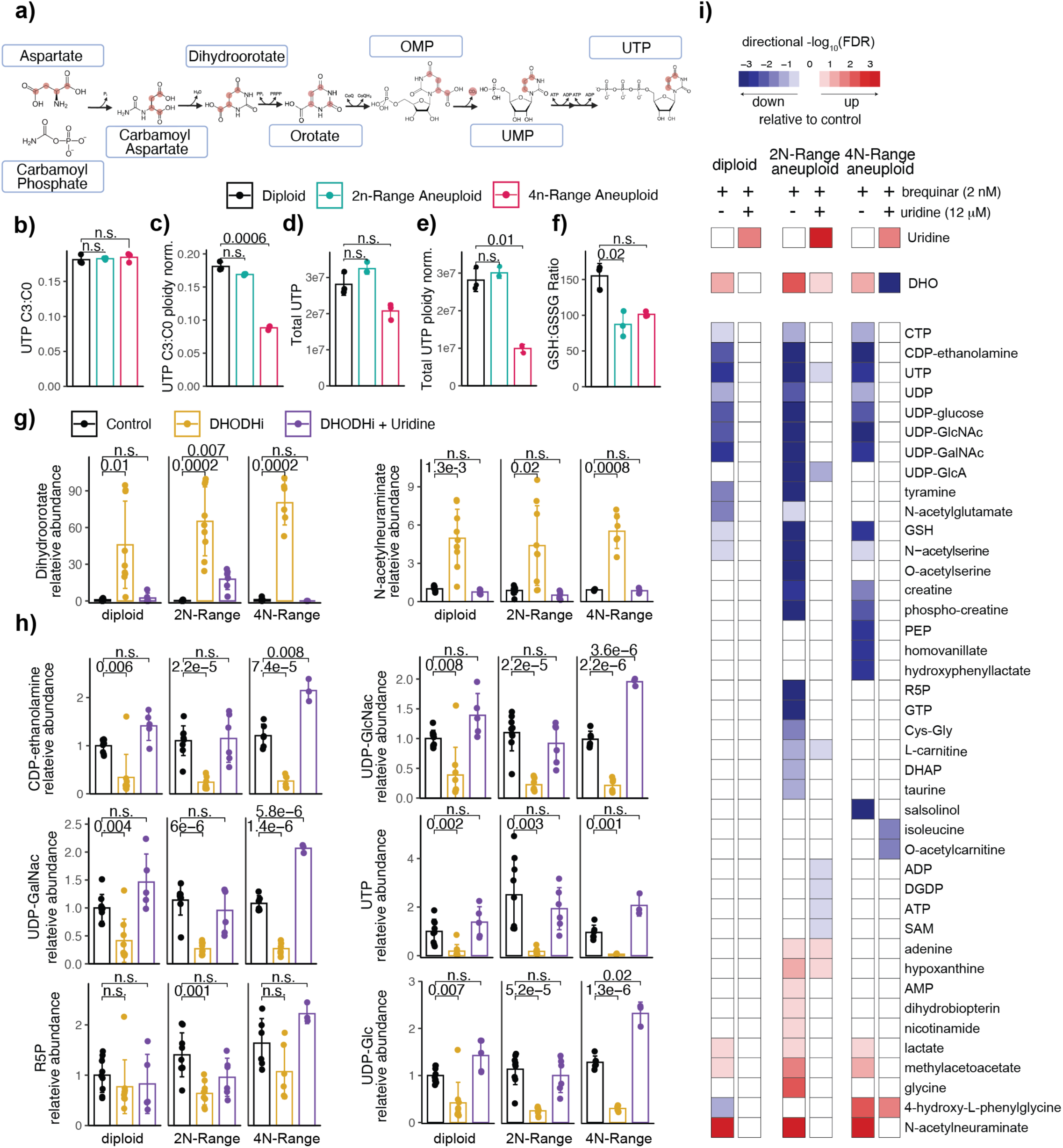
Pyrimidine synthesis inhibition exacerbates nucleotide insufficiency in net-gain aneuploid cells. **a,** Diagram depicting the labeled-carbon aspartate tracing through the pyrimidine biosynthesis pathway. **b,** Bar graphs showing the ratio of labeled UTP to unlabeled UTP from the aspartate tracing experiment in diploid, 2N-range-aneuploids, and 4N-range-aneuploids. **c,** Ratio of labeled UTP to unlabeled UTP normalized for total chromosomal content relative to the diploid (46 chromosome) baseline. **d,** Total UTP (labeled + unlabeled) levels in each cell type, per-cell-normalized. **e,** Total UTP (labeled + unlabeled) levels in each cell type, normalized for total chromosomal content. **f,** Ratio of reduced (GSH) to oxidized (GSSG) glutathione in each cell type. **g, h,** Bar graphs showing the relative abundance of two metabolites that increase significantly with low dose DHODH inhibition (2nM brequinar), Dihydroorotate and N-acetylneuraminate **(g)**, as well as several pyrimidine-containing metabolites that decrease **(h)** in diploid, 2N-range aneuploid, and 4N-range aneuploid cells. P-values were calculated using T tests corrected for multiple testing. **i,** Heatmap summary of LC/MS-MS-derived metabolomic profiles of diploid, 2n-range aneuploid, and 4n-range aneuploid cells treated with DHODHi, with or without uridine rescue, as compared to control conditions for each line. Conditions shown are: untreated, 2nM brequinar-treated, and 2 nM brequinar + 12 μM-uridine-treated. FDR values were calculated from edgeR-based analysis corrected for multiple hypothesis testing.

Underlying insufficiency in *de novo* pyrimidine flux relative to nucleodide demand suggests a mechanism for sensitiviy of aneuploid HMECs to DHODH disruption, which would exacerbate this pre-existing insufficiency. We thus examined the underlying metabolic disruptions induced by low-dose DHODHi (2nM brequinar) in aneuploid and diploid cells using LC-MS/MS metabolomics. As expected, a buildup of the DHODH substrate, dihydroorotate (DHO), was observed with DHODHi, validating the on-target nature of the drug (**Fig. 3 g**). Diploid cells displayed minimal metabolomic changes associated with low-dose DHODHi, primarily a decrease in pyrimidine species, which were restored to baseline levels with uridine supplementation (**Fig. 3 g-i and Fig. S3**). However, 2N- and 4N-range net-gain aneuploid cells exhibit greater metabolomic disruption with DHODHi beyond just decrease in pyrimidine species (which were also more significantly decreased than in diploids) (**Fig. 3 g-i and Fig. S3**). Furthermore, in aneuploid HMECs, uridine supplementation rescues most (but not all) of the metabolic disruption caused by DHODHi (**Fig. 3 g-i and Fig. S3**).

### Both oxidative phosphorylation and glycolytic rates are increased in net-gain aneuploid HMECs

Since we also observed increased dependence of net-gain aneuploid cells on mitochondrial pathways including oxidative phosphorylation, and metrics of redox stress like GSH::GSSG ratio were also apparent in aneuploid HMECs (**Fig. 3 f**), we asked whether baseline bioenergetic states of aneuploid and diploid cells were altered. We found that aneuploidy increases both oxygen consumption and proton efflux rates to a degree scaling with overall ploidy, indicative of increased oxidative phosphorylation and glycolytic flux, respectively (**Fig. 4 a-f and Fig. S4 a, b**). We next assessed total mitochondrial content relative to DNA ploidy across a broad range of aneuploidies in HMECs and found an increase in estimated mitochondrial DNA in 4N-range aneuploids (even relative to ploidy), along with an increase in both nuclear-encoded and mitochondria-encoded mitochondrial gene expression (**Fig. 4 g-k)**. Thus, mammary epithelial cells with higher chromosomal ploidies tend to increase mitochondrial content and mitochondrial gene expression and display higher rates of oxidative phosphorylation. The increased flux through oxidative phosphorylation and glycolysis observed in net-gain aneuploids may serve to compensate for increased energy expenditure, since ratios of ATP::AMP are largely equivalent in aneuploids and diploids (**Fig. S4 c**).

**FIGURE 4:**
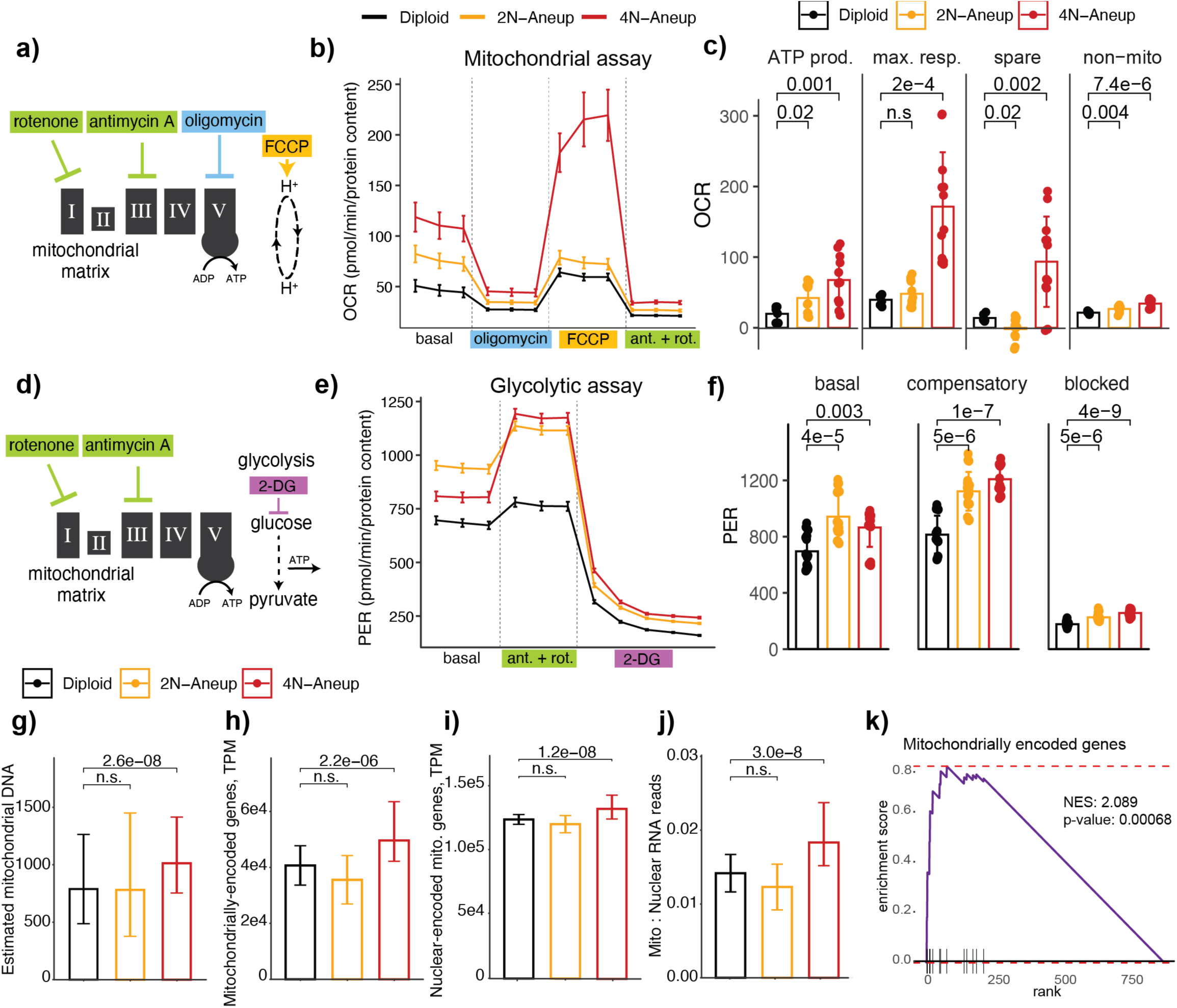
Net-gain aneuploid HMECs exhibit increased mitochondrially-linked ATP production and increased glycolytic rate. **a,** Diagram depicting the point of action of the drugs used in the Mitochondrial Stress Test Assay. **b,** Oxygen consumption rate (OCR) of diploid, 2n-range aneuploid, and 4n-range aneuploid cells after treatment with each of the drugs in the assay. **c,** Bar graphs summarizing average baseline ATP production, maximal respiration, spare capacity, and non-mitochondrial respiration in each cell type. P-values derived from T tests, corrected for multiple hypotheses. **d,** Diagram depicting the point of action of the drugs used in the Glycolytic Rate Test. **e,** Proton efflux rate (PER) of diploid, 2n-range aneuploid, and 4n-range aneuploid cells after treatment with each of the drugs in the assay. **f,** Bar graphs summarizing average normalized PER levels during basal, compensatory, and blocked phases of the assay in diploid, 2N-range aneuploid, and 4N-range aneuploid cells. P-values derived from T tests, corrected for multiple hypotheses. **g-j,** Bar graphs showing the estimated number of mitochondrial DNA copies relative to nuclear DNA content **(g)**, transcripts per million (TPM) of mitochondrially-encoded genes **(h)**, TPM of nuclear-encoded mitochondrial genes **(i)**, and the ratio of mitochondrial to nuclear RNA reads **(j)** in diploid, 2N-range aneuploid, and 4N-range aneuploid cells. **k,** Mitochondrially-encoded gene set enrichment in 4N-range aneuploid HMECs compared to diploids. Enrichment score and P-value from the analysis are shown.

### Net gain aneuploid HMECs activate p53 and are sensitive to DNA-damaging drugs combined with DHODHi

To assess the relationship between nucleotide pool insufficiency and stress signaling in aneuploidy, we performed RNA-seq in representative aneuploid and diploid HMEC lines under steady state, mild nucleotide inhibition with DHODHi, or DHODHi plus UMP supplementation conditions. We validated that copy number-specific gene expression changes could be observed in this gene set, as expected (**Fig. S5 a-d**). Low-dose DHODHi (2 nM brequinar) resulted in decreased ribosomal gene expression universally across all cell types, albeit to more significant degrees in aneuploid cells (**Fig. 5 a, b**) Diploid transcriptomes were otherwise unperturbed by DHODHi, and the modest ribosomal gene downregulation was fully rescued by uridine salvage. 2N- and 4N-range net-gain aneuploid HMECs displayed more significant and widespread transcriptomic responses to DHODHi, including a robust activation of p53 signaling as indicated by canonical target upregulation (**Fig. 5 a, b**), as well as activation of lysosomal, cell adhesion, and NFκB target genes (**Fig. S5 e**). Unlike in diploid cells, transcriptomic alterations in response to DHODHi were not fully restored to baseline with uridine supplementation in aneuploid HMECs (**Fig. 5 a**).

**FIGURE 5:**
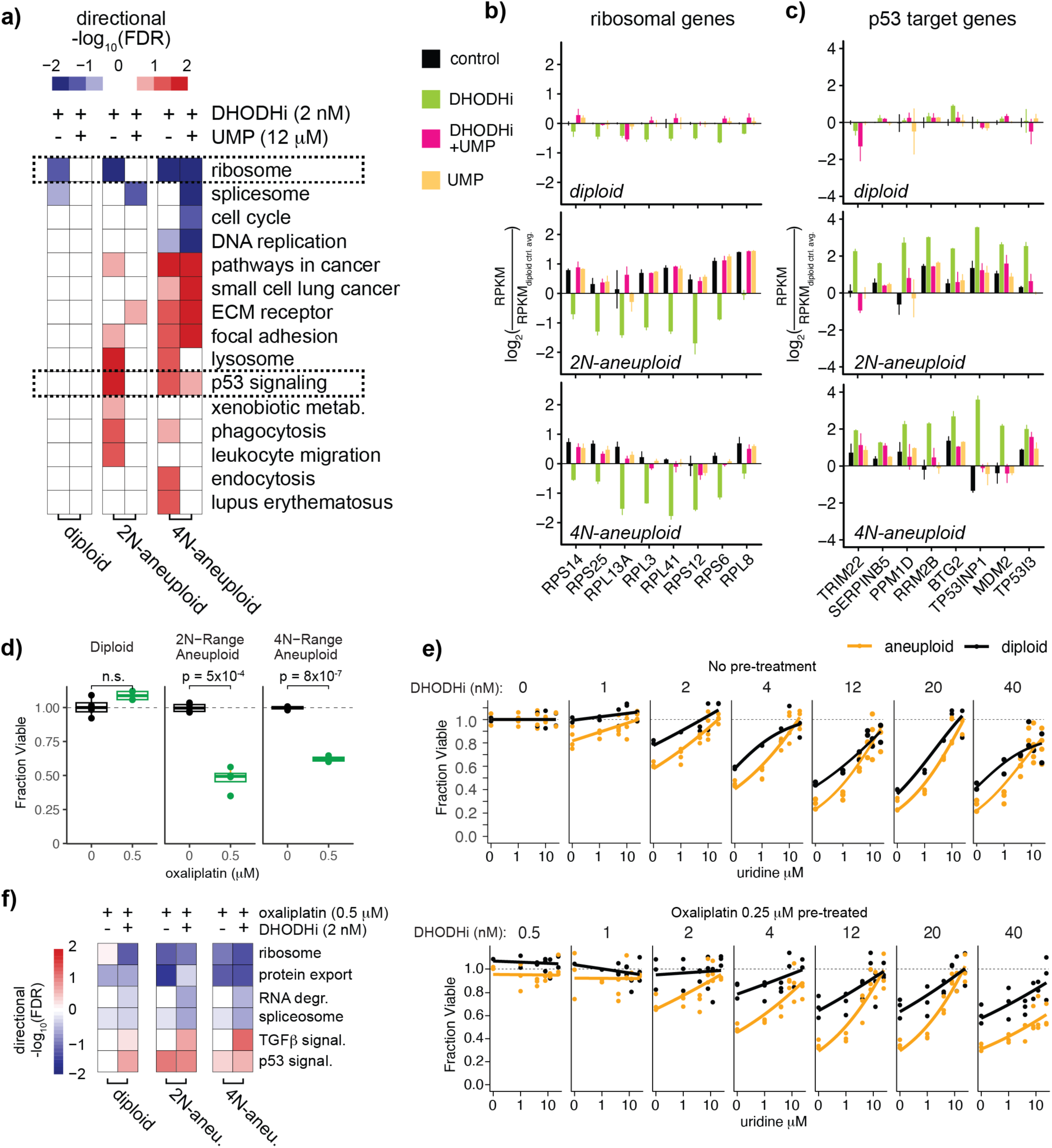
Nucleotide pool stress signaling is activated by low-dose DHODHi and DNA damaging agents in net-gain aneuploid HMECs. **a,** Heatmap summary of transcriptomic profiles of diploid, 2N-range aneuploid, and 4N-range aneuploid HMECs treated with the DHODH inhibitor brequinar or with a combination of brequinar and UMP, relative to control media conditions. Relative expression levels of ribosomal genes and p53 target genes are outlined with a dotted black box. **b, c,** Bar graphs showing relative reads per kilobase per million reads (RPKM) of several ribosomal genes **(b)** and p53 target genes **(c)** in diploid, 2N-range aneuploid, and 4N-range aneuploid cells treated with brequinar, brequinar and UMP, UMP, or under control media conditions. **d,** Cell viability of untreated diploid, 2N-range aneuploid, and 4N-range aneuploid HMECs treated with 0.5 μM oxaliplatin compared to control media conditions. P-values were calculated using T tests. **e,** Cell viability of diploid (shown in black) and 2N-range aneuploid HMECs (shown in yellow) that have either undergone no pre-treatment (top) or pre-treatment with 0.25 μM of oxaliplatin (bottom), and were subsequently challenged with brequinar ranging from 0-40 nM, with uridine ranging from 0-10 μM. **f,** Heatmap summary of transcriptomic profiles of diploid, 2N-range aneuploid, and 4N-range aneuploid HMECs treated with either oxaliplatin or with a combination of oxaliplatin and brequinar, as compared to control conditions.

We wondered whether aneuploidy-associated nucleotide pool stress may result in sensitivity to chemotherapeutic DNA-damaging agents, which would exacerbate replication stress and nucleotide pool demand to support DNA repair. We exposed cells to 0.5 μM oxaliplatin for 3 days and found that aneuploid cells were significantly more sensitive to chemotherapeutic challenge than diploid cells (**Fig. 5 d**). To test the combination of both DNA damaging and DHODHi drugs, we pre-treated cells with 0.25 μM oxaliplatin for 3 days to induce damage, then inhibited pyrimidine biosynthesis across a range of DHODHi (brequinar) doses and uridine rescue conditions. Pre-treatment with oxaliplatin exacerbated the differential proliferative effects of DHODHi in aneuploid cells compared to diploid cells across a range of salvage conditions, but especially when salvage is restricted by low uridine levels (**Fig. 5 e**). Additionally, 0.5 μM oxaliplatin treatment for 3 days resulted in a similar transcriptional response observed by DHODHi-induced nucleotide pool stress including downregulation of ribosomal gene expression and increased p53 signaling, but only in aneuploid cells, and this response was even more pronounced when treatments were combined **(Fig. 5 f)**.

### Net-gain aneuploid tumors and cancer cell lines display similar metabolic features

To determine whether aneuploidy-associated nucleotide insufficiency phenotypes were also present in human tumors and cancer cell lines, we assessed large-scale metabolomics, sequencing, and functional genomics datasets. A paired RNA-seq/metabolomics dataset of 108 breast carcinoma samples with matched normal tissue^52^ enabled us to infer copy number profiles of tumors (**Fig. S6 a**) and assess metabolic trends associated with net-gain tumors. We performed a similar analysis of 907 cancer cell lines (pan-subtype) with paired copy number and metabolomics data^53,54^. Several metabolic shifts that were observed in net-gain aneuploid HMECs relative to diploid HMECs at steady state, including increased TCA cycle intermediates (citrate), oxidized glutathione (GSSG), and phosphoenolpyruvate (PEP), were also observed in net-gain cancer cell lines and breast carcinoma samples (**Fig. 6 a-c and S6 b-d**). This may indicate increased TCA cycle/oxidative phosphorylation activity in addition to increased glycolysis rates associated with net chromosomal gain, as confirmed with bioenergetic assays in HMECs (**Fig. 4**).

**FIGURE 6:**
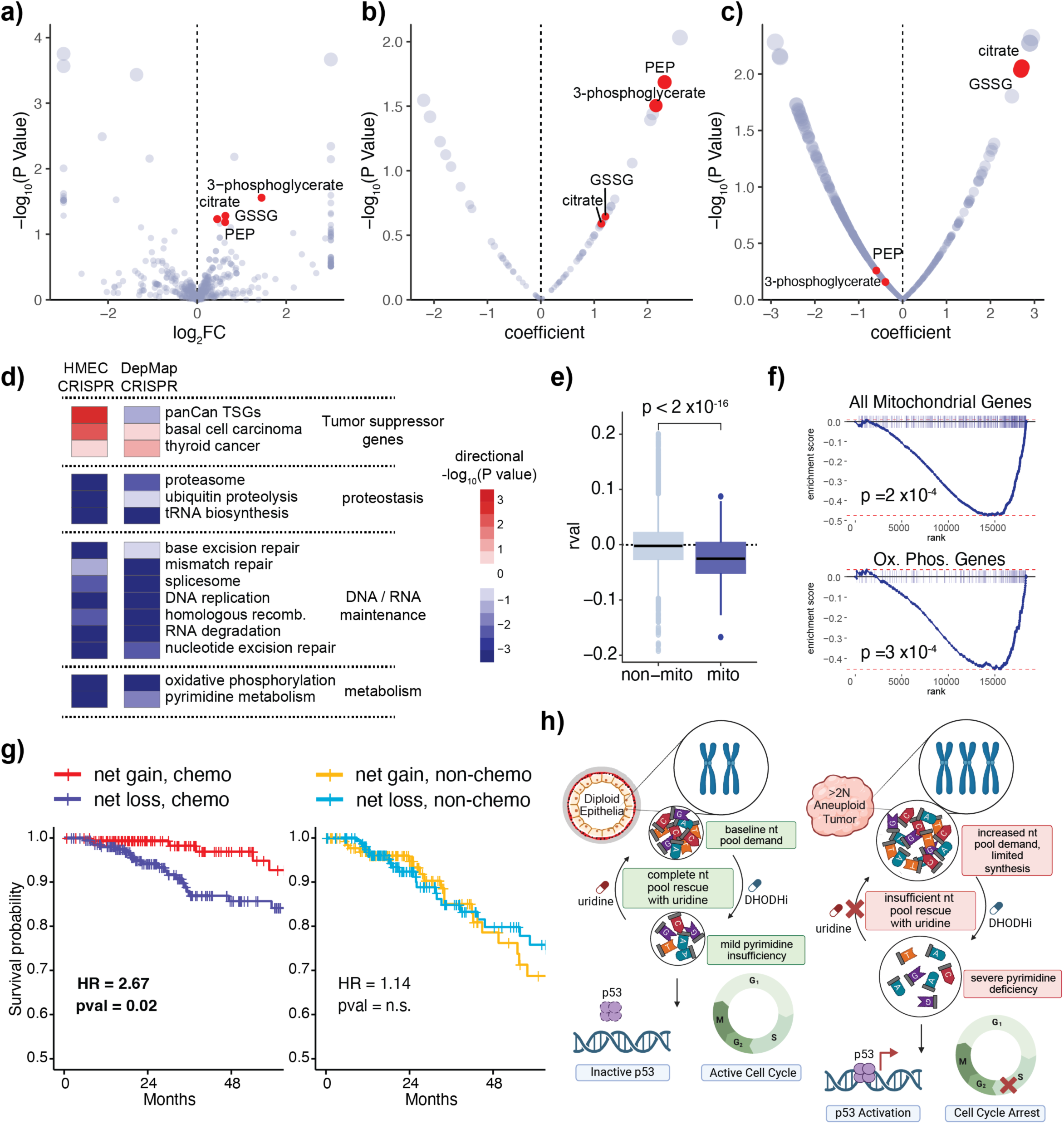
Net-gain aneuploidy is associated with metabolic phenotypes and is prognostic for response to DNA-damaging chemotherapy in cancer. **a,** Volcano plot showing metabolite log_2_ fold change (log_2_FC) (x-axis) and -log_10_(P-value) (y-axis) in net-gain aneuploid HMECs compared to diploid HMEC controls under steady state conditions. log_2_FC capped at minimum -3 and maximum 3. **b,** Volcano plot showing linear regression coefficient (x-axis) and corresponding -log_10_(P-value) of metabolite levels compared to net-gain aneuploidy levels across cancer cell lines. **c,** Volcano plot showing linear regression coefficient (x-axis) and corresponding -log_10_(P-value) of metabolite levels compared to net-gain aneuploidy levels across a cohort of human breast cancer samples (aneuploidy levels inferred from matched RNA-seq data). **d,** Heatmap of differentially essential gene sets associated with net-gain aneuploidy across DepMap cancer cell line CRISPR screen data (right column), shows a similar pattern to the aneuploidy epistasis profile of HMECs (left column). Gene set enrichment analysis (KEGG gene sets) was performed on a gene list ranked by the correlation of effect scores with net-gain aneuploidy levels. **e,** Gene effect scores of mitochondrially-localized genes are mostly negatively correlated with net-gain aneuploidy levels across cancer cell lines. **f,** Gene set enrichment plots showing all mitochondrially-localized genes (top) or just oxidative phosphorylation genes (bottom) using the gene ranking in **(d)**. **g,** Overall survival across a 5-year timeframe of breast cancer patients from the TCGA cohort with aneuploid tumors classified as either ‘net-gain’ or ‘net-loss’, treated with DNA-damaging/anti-nucleotide/anti-metabolite chemotherapeutics (left panel) or without chemotherapy (right panel). Hazard ratios and associated P-values were calculated from Cox proportional hazards regression models. **h,** Diagram depicting a net-gain aneuploidy nucleotide insufficiency model and putative therapeutic vulnerability. Increased nucleotide pool requirements render net-gain aneuploid cells sensitive to nucleotide pool stress caused by nucleotide synthesis inhibitors or DNA-damage. This nucleotide pool stress cannot be fully rescued by uridine salvage, and results in p53 activation and cell cycle arrest, whereas diploid cells can replicate efficiently using salvage alone.

To determine what genetic dependencies correlated with net-gain aneuploidy broadly across cancer cell lines, we assessed the DepMap CRISPR data^55^ along with chromosomal copy number data across 756 cancer cell lines. Ranking gene effects by their correlation to the degree of net-gain aneuploidy in cancer cell line genomes and performing gene set enrichment analysis revealed a similar pathway-level synthetic lethality profile compared to the HMEC aneuploidy synthetic lethality screens in this study (**Fig. 6 d** and **S6 e**). Top synthetic lethal pathways with net-gain aneuploidy in cancer cell lines included oxidative phosphorylation, pyrimidine metabolism, DNA repair pathways and proteostasis pathways (**Fig. 6 d and S6 e**). The essentiality of genes encoding mitochondrially localized proteins was generally correlated with the degree of net-gain aneuploidy across cancer cell lines (**Fig. 6 e, f**). A similar overall gene set enrichment profile was observed when analyzing net-gain aneuploidy in just breast cancer-derived cell lines (n = 33) (**Fig. S6 f**). This indicates that net gain aneuploidy universally confers sensitivity to disruption of nucleotide, DNA repair, and mitochondrial pathways, in both non-cancer-derived and cancer-derived cell line models.

### Breast cancer patients with net-gain aneuploid tumors have increased survival probability when treated with DNA-damaging chemotherapies

These combined functional genomics analyses suggest that nucleotide insufficiency in net-gain aneuploid tumors may represent a therapeutically exploitable vulnerability to drugs that affect the nucleotide pool and DNA replication, such as DNA-damaging chemotherapeutics (like platin-based therapies), nucleotide analogs (like 5-FU), or anti-metabolites (like methotrexate). Indeed, stratifying patients in the breast cancer TCGA cohort by aneuploidy (net gain vs. net loss) and treatment type (chemotherapy vs non-chemotherapy) revealed that patients with net-gain tumors treated with chemotherapy have a significantly higher survival probability than patients with net-loss tumors (HR = 2.67, p = 0.02) (**Fig. 6 g**). This difference in survival was not observed in patients who were treated with non-chemotherapeutic hormonal and/or targeted therapies (**Fig. 6 g**). The best overall survival was achieved in the net-gain aneuploidy + chemotherapy treatment group (95% 5-year survival rate) (**Fig. 6 g**). This suggests that aneuploidy status could be used in therapeutic decision-making and patient stratification, and that patients with net-gain aneuploid tumors could benefit significantly from receiving DNA-damaging/nucleotide analog/anti-metabolite chemotherapies. Incorporating biosynthesis inhibitors like DHODHi into treatment regimens may provide additional benefit by exploiting the inability of net-gain aneuploid cells to proliferate using pyrimidine salvage alone (**Fig. 6 h**). Minimal uridine supplementation could be a way to buffer systemic toxicity but still maintain effectiveness in aneuploid cell growth restriction (**Fig. 6 h**).

## Discussion

In this study we performed genome-wide CRISPR knockout screens in human mammary epithelial cells with net-gain aneuploidy compared to isogenic diploid cells. In addition to known aneuploidy stress-related susceptibilities like ubiquitin-mediated protein homeostasis, we identified mitochondrial oxidative phosphorylation, TCA cycle, and de novo pyrimidine biosynthesis enzymes as top-scoring dropout hits in the screen. Additionally, we observed increased enrichment of sgRNAs targeting components of the p53 pathway in aneuploid cells compared to diploids, indicating that aneuploid cells derive greater benefit from loss of commonly mutated tumor suppressors. Aneuploidy has been shown to activate TP53 signaling in multiple cell types^11,18,25,56^ including HMECs^3^ (**Fig. 5a-b**), and loss of TP53 via mutation licenses aneuploidy *in vitro*^28^ and is associated with increased aneuploidy in human tumors^34,35^. Replication stress has been previously proposed as a mechanism linking aneuploidy and p53 activation^18^, however the root causes of aneuploidy-associated replication stress are not well understood. Here, we identify nucleotide pool insufficiency as a metabolic vulnerability of net-gain aneuploid HMECs, which is associated with p53 activation and exploitable with pyrimidine synthesis inhibitors.

DHODH appears to be a key node linking increased reliance on mitochondria and nucleotide metabolism in aneuploid cells. DHODH is the sole mitochondrial-resident enzyme of the *de novo* pyrimidine biosynthesis pathway, which reduces ubiquinone in the dehydrogenation of dihydroorotate to orotate. While we observe significantly increased ATP production via oxidative phosphorylation, as well as increased glycolysis rates, in net-gain aneuploids, we do not observe increased flux through *de novo* pyrimidine biosynthesis, perhaps revealing why it is the top vulnerability of aneuploid HMECs. Knocking out DHODH with CRISPR revealed that, while diploid HMECs could proliferate using solely *de novo* pyrimidine synthesis or salvage (with the ability to switch seamlessly between the two), aneuploid HMEC proliferation was significantly impaired under the salvage-only condition. This indicates a constitutive reliance on *de novo* synthesis in mammary epithelial cells with extra chromosomes even when nucleotides are plentifully available for salvage, perhaps due to a fixed upper limit on import or salvage capacity (adding high levels of uridine couldn’t override the deficiency).

Mild impairment of DHODH function with low-dose brequinar treatment exacerbated the underlying p53 activation phenotype associated with aneuploidy in HMECs and led to global disruption of the metabolome, including depletion of almost all pyrimidine-based species (CTP, CDP, UTP, UDP, and UDP-conjugated sugars). Sensitivity to DHODH inhibition in net-gain aneuploid cells may ultimately be due to a mismatch between nucleotide synthesis rates and nucleotide demand. Interestingly, ribosomal gene expression is increased at baseline in aneuploid cells relative to diploids, then decreases dramatically in response to DHODHi (**Fig. 5 b**, black bars). Ribosome biogenesis could be expected to compete with DNA replication for nucleotide pools given that the total rRNA content in a human cell (∼8-16 pg) is greater than the total amount of genomic DNA (∼6 pg)^57^, thus the interplay between these major consumers of pyrimidines could be an intersection of vulnerability in net-gain aneuploid cells. p53 activation occurs in response to both rRNA/ribosome biogenesis disruption and replication stress^58^, again highlighting the central role of p53 in aneuploidy stress sensing and response.

Combination treatment of cells with brequinar and oxaliplatin showed that treatment with DNA-damagers increases cell sensitivity to DHODH inhibition, particularly in aneuploid cells. This data suggests the potential for adjuvant therapies utilizing smaller doses of nucleotide inhibitors in combination with other clinically available DNA damaging chemotherapies, which have been shown to increase reliance on *de novo* pyrimidine synthesis in breast cancer^59^. Additionally, physiological doses of uridine are more efficient in returning diploid cells to baseline transcriptomic and metabolomic conditions as compared to aneuploid cells, therefore allowing diploid cells to be successfully rescued from arrest in S-phase. This data suggests a potential for limited nucleotide supplementation to be used in conjunction with nucleotide inhibitors to titrate the toxicity of chemotherapies.

Altogether, we show that aneuploid cells with a net gain of chromosomes experience nucleotide pool stress due to an increased demand of nucleotides but insufficient compensatory flux through synthesis and/or salvage pathways. Furthermore, p53 activation in net-gain aneuploid mammary cells may result from intrinsic metabolic constraints on pyrimidine synthesis and salvage leading to replication stress, which can be further exacerbated by pyrimidine synthesis inhibitors or DNA damage. Nucleotide salvage should be further investigated as a targetable node, as has been explored in kidney cancers with FH mutation and subsequent reliance on purine salvage^60^. Since many tissues can access both synthesis and salvage of nucleotides^61^, further exploration of the generalizability of our findings to net-gain aneuploidy in other tumors types is warranted. Survival data across a large cohort of breast cancer patients indicate that net gain aneuploidy could be utilized as a biomarker to identify patients who are likely to respond to DNA-damaging and anti-metabolite/anti-nucleotide treatments.

## Methods

### Cell Culture

The hTERT–HMEC cell line was immortalized previously^62^ from primary HMECs purchased from ATCC (PCS-600-010). Low-passage hTERT–HMECs were grown in Lonza HMEC medium with bovine pituitary extract and growth supplements. Cells are passaged once they reach ∼90% confluency using 5 mL of 0.05% Trypsin. We have previously confirmed the identity of this cell line using DNA-seq and RNA-seq analysis^3^. Human embryonic kidney (HEK) 293T cells used for lentiviral production were cultured in DMEM supplemented with 10% FBS and 1% Pen-Strep.

### Genome-wide CRISPR screens

A genome-wide CRISPR library targeting approximately 18,000 genes with 5 sgRNAs per gene was utilized in the lentiCRISPR v2 backbone. The library was packaged along with lentiviral packaging components (Tat, Rev, Gagpol, and Vsvg) into lentiviral particles using lipofectamine transfection of HEK 293T cells in lentiviral packaging media (Opti-MEM reduced serum media with 5% FBS and 1x GlutaMAX supplement). Lentivirus was harvested and concentrated using Lenti-X Concentrator. Target HMECs were infected with the library at a multiplicity of infection (MOI) of 0.3 to ensure single viral integration per cell. A representation of 500x was maintained throughout the experiment to ensure robust screening coverage. Following infection, HMECs were selected with puromycin at a concentration of 2 μg/mL for 2 days to enrich for cells that were successfully transduced with the lentiviral constructs. Post-selection, cells were allowed to expand in culture and were harvested at two time points: immediately after puromycin selection (timepoint 0) and after the cells had undergone approximately 6 population doublings (PD6). Genomic DNA was isolated from the collected cell pellets after lysis and proteinase K treatment using the phenol-chloroform extraction method. The regions containing the integrated sgRNA sequences were amplified by PCR to generate amplicons for sequencing. Amplicon sequencing was performed on the Illumina NextSeq 550 platform to quantify the abundance of each sgRNA construct in the cell populations. Sequencing reads were aligned to the reference library using Burrows-Wheeler Aligner (BWA)^63^. The differential depletion or enrichment of sgRNAs between timepoint 0 and PD6 was analyzed using the limma-voom pipeline in conjunction with edgeR^43,44^ and the camera^64^ function. Gene set enrichment analysis was performed using the fgsea^65^ package to identify significantly enriched pathways or gene sets.

### Generating DHODH-/- cell lines with CRISPR

The lentiCRISPR v2 backbone^66^ was digested with the BsmBI restriction enzyme, and the cut vector was isolated through a gel extraction. sgRNA guides targeting DHODH were then ligated into the digested lentiCRISPR v2 backbone and packaged into lentivirus though transfection of HEK 293T cells along with lentiviral packaging components (Tat, Rev, Gagpol, and Vsvg) with Lipofectamine 3000 in lentiviral packaging media made of Opti-MEM reduced serum media with 5% FBS and 1x GlutaMAX supplement. Lentivirus was then harvested and passed through a 0.45 μm filter and used to transduce HMECs. Transduced cells were then selected with 2 μg/mL of puromycin for 2 days and recovered in HMEC media as described above, supplemented with 20 μM of uridine. Reduction in levels of DHODH protein was quantified through western blots. The sgRNA target sequence for the control AAVS1 locus is GGGGCCACTAGGGACAGGAT, and sgRNA target sequence for DHODH is GTGTTCGCTTCACCTCCCTG.

### Western Blots

Cell pellets of approximately 5 x 10^5 cells from each HMEC line were lysed in 250 μL 2x RIPA buffer plus protease inhibitor cocktail. Lysates were vortexed and spun down, and protein concentrations were determined by bicinchoninic acid protein assay (Pierce cat. no. 23227), then equal amounts of protein were mixed with Pierce Lane Marker Reducing Sample Buffer and loaded onto 4–12% Bis-Tris gels, 1.5 mM, with 15 wells (Invitrogen cat. no. NP0336BOX). Gels were run in MOPS SDS buffer (Life Technologies cat. no. NP0001) and transferred to nitrocellulose (BioRad cat. no. 170-4158), blocked overnight in 3% BSA at 4 °C, then incubated overnight at 4 °C with DHODH antibody (DHODH (E9X8R) rabbit mAb, Cell Signaling cat. no. 26381S) at 1/1000 dilution in TBST buffer with 1% BSA, or with Vinculin antibody (Vinculin (EPR19579) rabbit mAb, Abcam cat. no. ab207440) at 1/1,000 dilution. Secondary antibody for all assays was goat anti-rabbit IgG (Abcam cat. no. ab205718), incubated at 1/10,000 dilution for 1 h at room temperature. Western blots were quantified using ImageJ^67^ v1.53a.

### Cell viability assays

HMEC cells of varying degrees of aneuploidy were seeded in the concentration of 5 × 10^5^ cells/ml in a final volume of 0.1 ml in 96-well flat-bottom microtiter plates. We used the fluorimetric resazurin reduction method (CellTiter-Blue; Promega) to evaluate the chemosensitivity of cells treated with brequinar and rescued with uridine (Sigma Chemical Co., St. Louis, MI, USA). These drugs were added in a range of 1–12 nM or 1–12 μM respectively and cells were treated for 48 hours. Fluorescence (537Ex/610Em) was determined using a luminometer (Tecan microplate reader). The percentage of viable cells was calculated using modeling technique.

### PI staining for total DNA content

Approximately 5 × 10^5^ cells per clone were fixed in 70% ethanol, then stored for up to 1 month at −20 °C. Fixed cells were spun down, fixative was removed and then cells were washed once in phosphate-buffered saline (PBS) and finally resuspended in 500 μl Thermo Fisher FxCycle PI/RNAse staining solution. After incubation in the dark for 30 min, cells were passed through a mesh filter sieve and analyzed by fluorescence-activated cell sorting (FACS) using 532-nm excitation with a 585/42-nm bandpass filter. An average of 1 × 10^4^ events were analyzed per clone, with data collected via BD FACSDiva software v.8.0 and processed using R packages flowCore^68^ and ggcyto^69^ to derive the average fluorescence of the G1 peak relative to that of diploid control cells processed simultaneously.

### RNA-seq library preparation and analysis

A total of 5 × 10^5^ cells from each cell line were plated in 10 cm plates and grown for 144 h. Cells were provided fresh media 3 h before collecting. Media was aspirated and cells were immediately lysed in dishes and total RNA was purified using Qiagen RNeasy kits. A quantity of 1 μg of total RNA was used for mRNA purification with the NEBNext Poly(A) mRNA Magnetic Isolation Module. NEBNext Ultra II Directional RNA Library Prep Kits for Illumina were used for RNA-seq library preparation. NEBNext Multiplex Oligos for Illumina were used for indexing during PCR amplification of the final libraries. Libraries were quantified by qPCR using the NEBNext Library Quant Kit for Illumina and multiplexed accordingly. RNA-seq reads were aligned to the human reference genome hg37 using BWA^63^. Following alignment, SAMtools^70^ was utilized to sort the aligned reads, which were then compiled into gene-level read counts using the subread featureCounts^71^ function. Gene-level differential expression was analyzed using the edgeR^43,44^ limma function and differential pathway analysis was performed with fgsea using KEGG^72^ gene sets.

### LC-MS/MS metabolomic profiling, aspartate tracing, and analysis Polar LC-MS Method

A Q-Exactive orbitrap mass spectrometer with an Ion Max source and HESI II probe attached to a Vanquish Horizon UHLPC system was used to measure polar metabolites. The LC-MS underwent weekly cleaning and calibration with positive and negative Pierce ESI Ion Calibration Calmix (Thermo Scientific). 2 μL of sample was injected into the machine and ran through a SeQuant ZIC-pHILIC 5 μm 150 x 2.1 mm analytical column (Sigma) with a 2.1 x 20 mm guard column (Sigma) attached to the front end. The column oven was set to 25°C and autosampler was set to 4°C. Buffer A was comprised of 20 mM ammonium carbonate (Sigma), 0.1% ammonium hydroxide (Sigma) in HPLC-grade water (Sigma) and Buffer B was compromised of 100% acetonitrile (Sigma). The liquid chromatography was set to a flow rate of 0.15 mL/min. First a linear gradient from 80% Buffer B to 20% Buffer B occurred over the course of 20 minutes followed by linear gradient from 20% Buffer B to 80% Buffer B for 0.5 minutes, followed by a hold at 80% Buffer B for 7.5min. The mass spectrometer was set to full scan (70-1000 m/z), polarity switching mode, with the spray voltage set to 4.0 kV, heated capillary to 350°C, and the HESI probe at 30 °C. The sheath gas flow was set at 10 units, auxiliary gas at 1 unit, and sweep gas flow at 1 unit. The resolution of scan was set to 70,000, AGC target to 1×10^6^, and maximum injection time at 20 msec. An additional scan between 220-700 m/z was used to enhance nucleotide detection in the negative mode as well with the maximum injection time set to 80 msec.

### Polar metabolite isolation from cultured cells

Media was aspirated from the plates and then the cells were washed with 1x PBS twice. The plate was then transferred to dry ice and 500 µL of 80% HPLC-grade methanol (Sigma) 20% HPLC-grade water (Sigma) was added to each well. The wells were placed in a -80 freezer to incubate for 15 minutes. The plates are taken out of the freezer one at a time and placed back on dry ice. The cells were then scraped and transferred to a new tube. Each well was washed with an additional 300 µL of 80% HPLC-grade methanol (Sigma) 20% HPLC-grade water (Sigma) and collected into the same tube as the initial lysis. The samples were then vortexed at 4°C for 10 minutes and centrifuged at 21,300 x g for 10 minutes at 4° C. Supernatants were transferred to a new tube and dried down in a Refrigerated CentriVap Benchtop Vacuum Concentrator connected to a CentriVap-105 Cold Trap (Labconco). After being dried down, pellets were stored in a -20° C freezer until ready to run on the polar LC-MS method.

### ^13^C_4_-aspartate tracing *in vitro*

Cells were seeded in Lonza HMEC medium 48 hours prior to tracing so that wells reached 75% confluence at time of experiment. 6 hours prior to metabolite isolation, the cells were treated with Lonza HMEC medium containing 10 mM ^13^C_4_-aspartate (Sigma) and the pH adjusted to 7.4 with the relevant treatments. After the 6-hour incubation period, polar metabolites were then isolated as described above and ran on the polar LC-MS method.

### LC-MS Data analysis

Metabolomics data were analyzed using TraceFinder 5.1 (ThermoFisher). Peaks were integrated using a strict 5 ppm mass tolerance and attention to the retention times as determined by purified standards of the respective metabolites. ^13^C and ^15^N-isotopologues were integrated with the same retention time as the ^12^C and ^14^N-isotopologues. All stable isotope tracing data underwent natural abundance correction using IsoCorrectoR^73^.

### Seahorse

Mitochondrial respiration and glycolytic rate of HMEC cells was measured as their oxygen consumption rate (OCR) and extracellular acidification rate (ECAR), respectively, using an oxygen-controlled XFe96 extracellular flux analyzer (Seahorse Bioscience). Diploid, 2N-range cells, and 4N-range cells were seeded in 12 replicates at 1×10^5^, cells per well, 2×10^5^ cells per well, and 2.5×10^5^ cells per well respectively in 80 μL of Lonza media into XFe96 cell culture microplates (Agilent) 24 hours prior to the experiment. Concurrently, the XFp sensor cartridge (Agilent) was hydrated by adding 200 μL of XF Calibrant (Agilent) and left in a CO2-free incubator overnight. An hour before the experiment, the Lonza media in the wells were replaced with 180 μL of Seahorse XF RPMI medium (Agilent) supplemented with 2 mM L-glutamine, 1 mM sodium pyruvate and 10 mM D-glucose. After 1 h incubation in a CO_2_-free incubator at 37 °C, glycolytic rate and mitochondrial stress tests were performed. Oxidative phosphorylation was assessed as follows utilizing the oxygen consumption rate (OCR). Initially, the basal OCR was measured, followed by the addition of 2.5 μM Oligomycin (Agilent), which inhibits complex 5 of the ETC and consequently decreases electron flow through the ETC. This allows for measurement of the ATP-linked respiration. Following this measurement, 2 μM of Carbonyl-cyanide-4 (triflouoromethoxy) phenylhydrazone (FCCP) (Agilent) was added, which is responsible for collapsing the proton gradient and disrupting the mitochondrial membrane potential, allowing for free flow of electrons through the ETC and measurement of the maximum mitochondrial respiration rate. The final injection includes a combination of rotenone and antimycin A, which block complex 1 and complex 3, respectively. This results in complete inhibition of mitochondrial respiration, allowing for the non-mitochondrial respiration to be measured. Glycolytic rate was assessed by first measuring the basal proton efflux rate (PER). In this assay, the first injection included a 0.5 μM combination of rotenone and antimycin A to block mitochondrial respiration. The second injection is 50 mM of 2-deoxy-D-glucose (2-DG) is then added, which competitively binds glucose-hexokinase and consequently inhibits glycolysis. The compensatory glycolytic rate was then measured by subtracting the PER after 2-DG injection from the PER after rotenone/antimycin A injection. Measurements were normalized using protein concentration obtained through the standardized protocol provided via the Thermo Scientific Pierce BCA Protein Assay kit.

### Nuclear:Mitochondrial DNA content analysis

Previously acquired DNA sequencing data (Watson et al, 2024) was used to estimate mitochondrial DNA numbers relative to nuclear DNA content using fastMitoCalc^74^.

### DepMap CRISPR screen analysis relative to copy number

Cancer cell line gene-level copy number information was used to rank cell lines by total net gain aneuploidy using a cutoff of log_2_(copy number ratio + 1) < 0.8 for losses and > 1.16 for gains. DepMap CRISPR screen data gene effect scores were correlated to net-gain aneuploidy levels across cell lines using linear regression analysis. Gene set enrichment analysis^75^ utilizing the KEGG gene sets^72^ was used via the fgsea^65^ package in R to identify gene sets that were epistatic with net-gain aneuploidy. To best compare with our screens, which were nucleotide supplement-free, we eliminated cancer cell lines from the analysis that were screened in media likely to contain nucleotide supplement (like F12, and medium 199) or for which media information was not available.

### Metabolomics analysis relative to copy number in cancer cell lines and tumors

Gene-level copy number data was used to determine net gain minus loss values in each cancer cell line^54^, using a cutoff log_2_(copy number ratio + 1) < 0.8 for losses and > 1.16 for gains. Gain minus loss values were then compared to metabolite levels^53^ across cell lines via linear regression analysis, using the lm function in R.

For human tumor analysis, copy number was estimated across the genome from RNA-seq data for each tumor sample using the CreateInfercnvObject function in the inferCNV package in R. Normal (non-cancerous tissue) samples from the same study were utilized as a reference group. Modified (normalized) expression was used to call gains or losses (> 1.02 or < 0.98, respectively) in genes likely attributable to copy number. Frequent breast cancer-associated events like +1q, -8p, +8q, and +16p could be observed in the inferCNV analysis, validating this methodology to detect cancer-associated CNAs from bulk RNA-seq data. Copy number calls from inferCNV were used to calculate gain minus loss values for each tumor, which were then compared to metabolite levels via linear regression analysis, using the lm function in R.

### Breast cancer survival analysis relative to copy number

Breast cancer copy number segment mean files generated by the TCGA Research Network (https://www.cancer.gov/tcga) were corrected for purity^76^ and arm-level gain/loss calls were made based on purity-corrected log_2_(copy number ratio) < -0.41 for losses and > 0.32 for gains, with least 50% of the arm affected. Gain minus loss values were calculated by subtracting the sum of total genomic Mb affected by chromosomal loss from the sum of total genomic Mb affected by chromosomal gain. Net-gain tumors were annotated as those with gain minus loss values > 0; net-loss tumors < 0. Patients receiving DNA-damaging, anti-folate, and anti-nucleotide chemotherapies were identified by the following search strings in the clinical datasets: “platin”, “dox”, “uracil”, “rubicin”, “mycin”, “phosphamid”, “citabine”, “cytoxan”, “trexate”, “ac”, “capecetabine”, “5-fu”, “metotreksat”, “mitoxantrone”, “cytoxen”, “gemzar”, “tc”, “tch”, “xeloda”. The survival (version 2.11-4)^77^ package in R was used for breast cancer patient survival analysis, utilizing the Surv() and survfit() functions. Survival data was plotted using the ggsurvplot function from the survminer (version 0.4.9)^78^ package in R. The coxph function in the survival package was used to fit a Cox proportional hazard regression model comparing outcomes in patients with net gain vs net loss tumors, in chemo- and non chemo-treated groups.

## Data Availability

All sequencing data is available on the NCBI SRA database, Bioproject ID PRJNA1165704.

## Acknowledgements

We would like to thank the Harvard Biopolymers Core for assistance with NextGen sequencing. E.V.W is supported by the Breast Cancer Alliance Young Investigator Award. This work was supported in part by an NIH grant R01CA234600 to S.J.E and the Harvard Ludwig Center, and the SPECIFICANCER Team funded by Cancer Research UK.

## Author Contributions

E.V.W. and S.J.E. designed the study. R.Y.M., A.N.K., F.N.K., A.F., T.T., A.C.L, J.A.H., and A.M.R.N. performed all experiments, with guidance from E.V.W., J.B.S., D.A.G., and S.J.E.. Data analysis was performed by R.Y.M., A.N.K., and F.N.K. under guidance from E.V.W.. The manuscript was written by R.Y.M., A.N.K., S.J.E., and E.V.W., with contributions from all other authors.

---

**SUPPLEMENT FIGURE 1:**
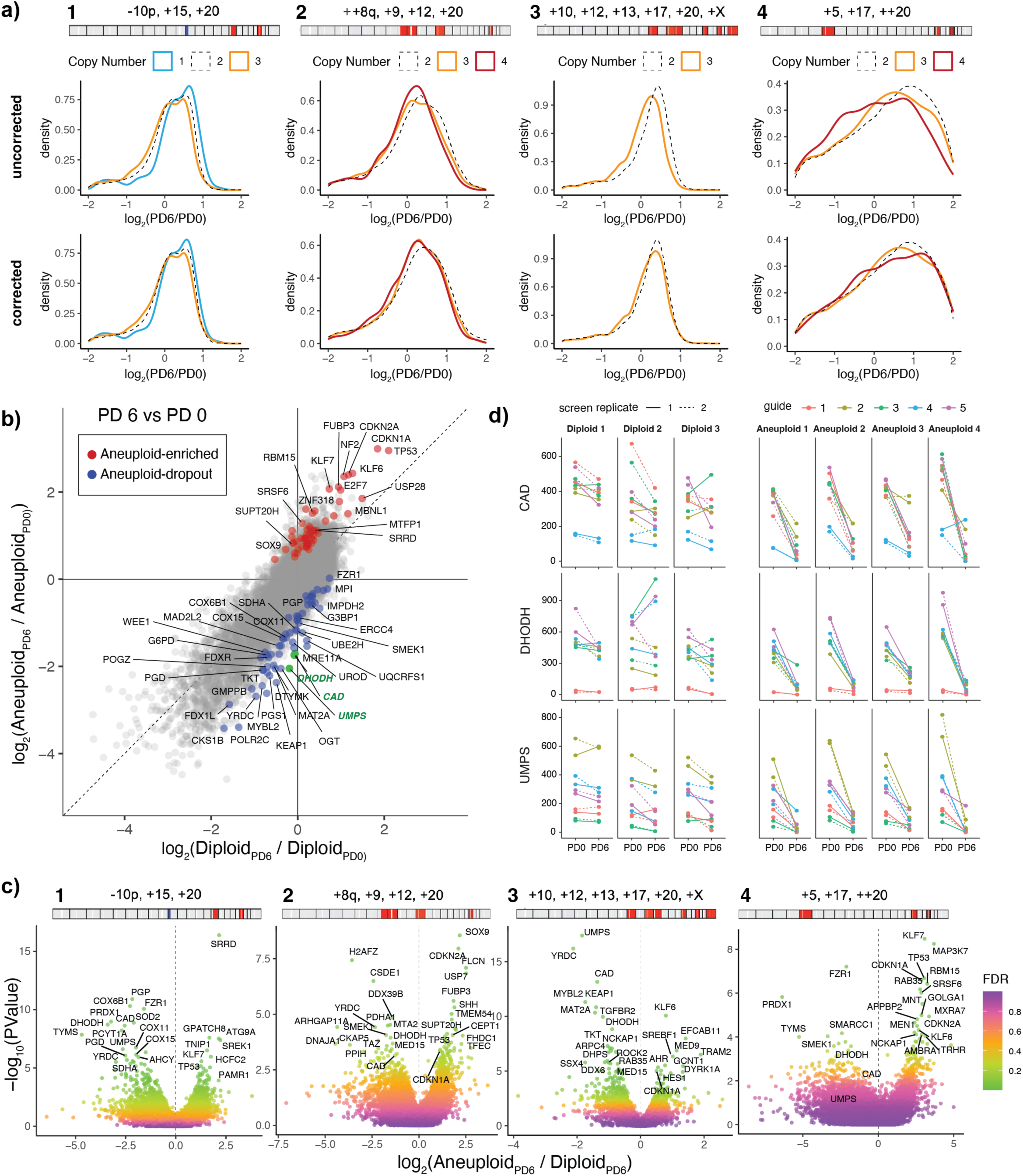
Genome-wide CRISPR knockout screens in aneuploid and diploid HMECs. **a,** Distribution plots showing uncorrected (top) and copy number-corrected (bottom) CRISPR screen log_2_(PD6/PD0) gene-level analysis, colored by gene ploidy level for each screen. Copy number profiles of aneuploid HMECs used in each screen are shown as colored bars above distribution plots. As previously reported^44^, genes on gained chromosomes tend to exhibit more dropout, and genes on lost chromosomes tend to exhibit less dropout, due to the dosage of CRISPR-mediated DNA cutting. **b,** Gene-level dropout and enrichment comparing PD 6 (End) sgRNA levels to PD 0 (Start) sgRNA levels in diploid (x-axis) and aneuploid (y-axis) CRISPR screens. Genes that are differentially depleted (blue) and enriched (red) in aneuploid HMECs relative to diploid HMECs at PD 6 are shown. Pyrimidine biosynthesis genes CAD, DHODH, and UMPS are colored in green. **c,** Differential dropout/enrichment of gene-targeting sgRNAs in aneuploid clones at PD 6 compared to their isogenic diploid clones at PD 6. Bars indicating the karyotype profiles of each of the aneuploid clones are shown at the top of the figure. **d,** Individual sgRNA counts targeting DHODH, UMPS, and CAD in cell populations at PD 0 and PD 6 in each replicate screen diploid (left) and aneuploid (right) HMEC lines.

**SUPPLEMENT FIGURE 2:**
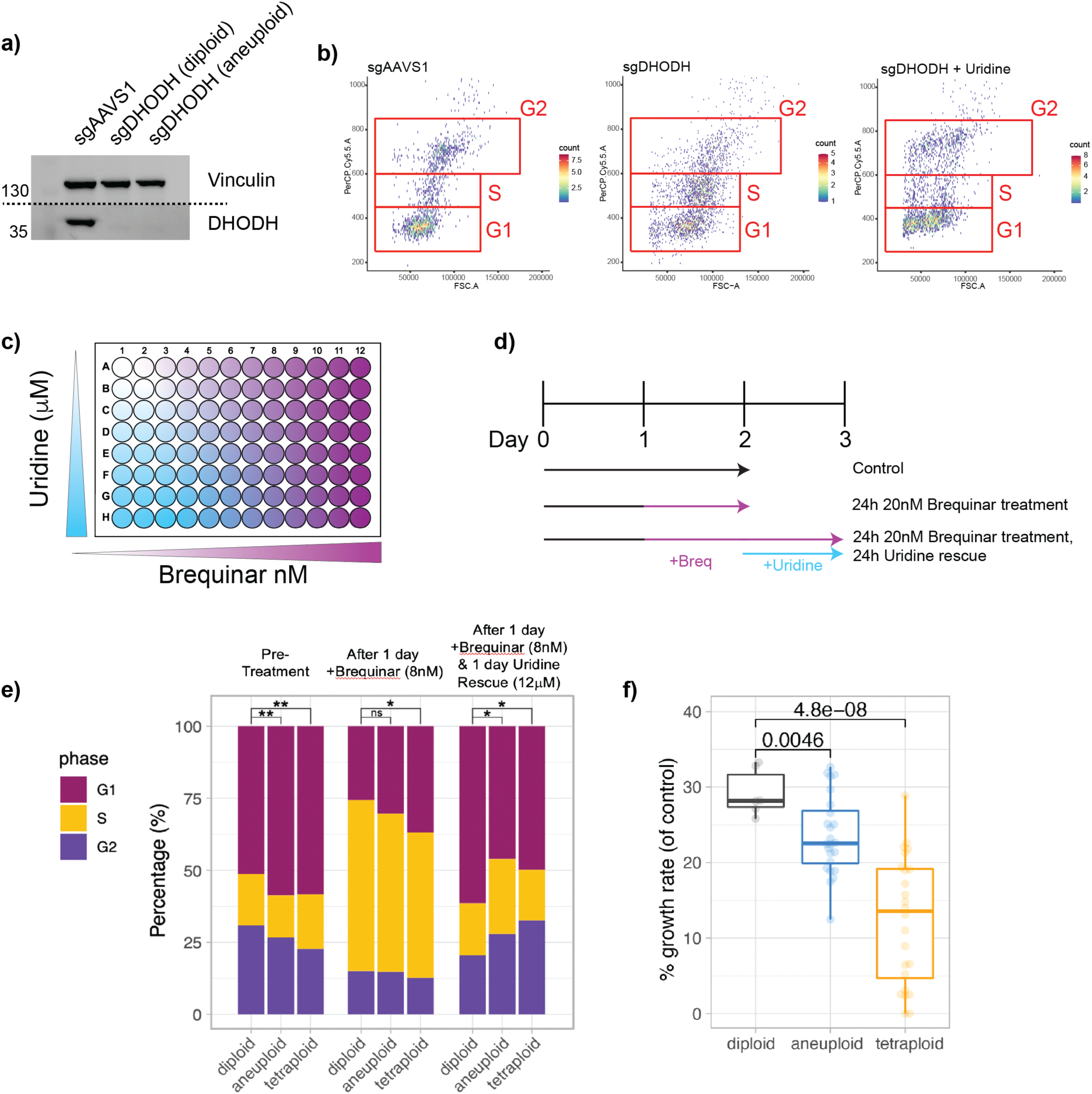
Net-gain aneuploid HMECs are more sensitive to acute or prolonged disruption of pyrimidine biosynthesis. **a,** Western blot showing complete knockout of DHODH in aneuploid and diploid cells exposed to DHODH-targeting sgRNA compared to an AAVS1-targeting control (in a diploid). DHODH knockout cells are maintained in uridine-supplemented media. **b,** Flow cytometry data showing forward scatter as a proxy for cell size on the x-axis and propidium iodide staining fluorescence to quantify total DNA content on the y-axis. Gates indicate cells in G1, S, and G2 phases of the cell cycle. **c,** Diagram showing the layout for the DHODHi (brequinar) + uridine supplementation matrix-style experiments in Fig XX. **d,** Diagram showing the dosing schedule for control conditions (black), acute brequinar treatment (purple) and a combination of acute brequinar treatment (purple) and uridine (blue) for experiments shown in subsequent panels. **e,** Percentage of diploid, 2n-range aneuploid, and 4n-range aneuploid cells in G1, S, and G2 phase during pre-treatment, under experimental conditions described in **(d)**. P-values were calculated from T tests corrected for multiple hypothesis testing, comparing percent of cells in S/G2 across cell types under each condition. **f,** The growth rate in diploid, 2N-range, and 4N-range aneuploid cells after DHODHi-mediated arrest and rescue for 24 hours with uridine, represented as the percent of control condition growth rate. P-values were calculated from T tests corrected for multiple hypothesis testing.

**SUPPLEMENT FIGURE 3:**
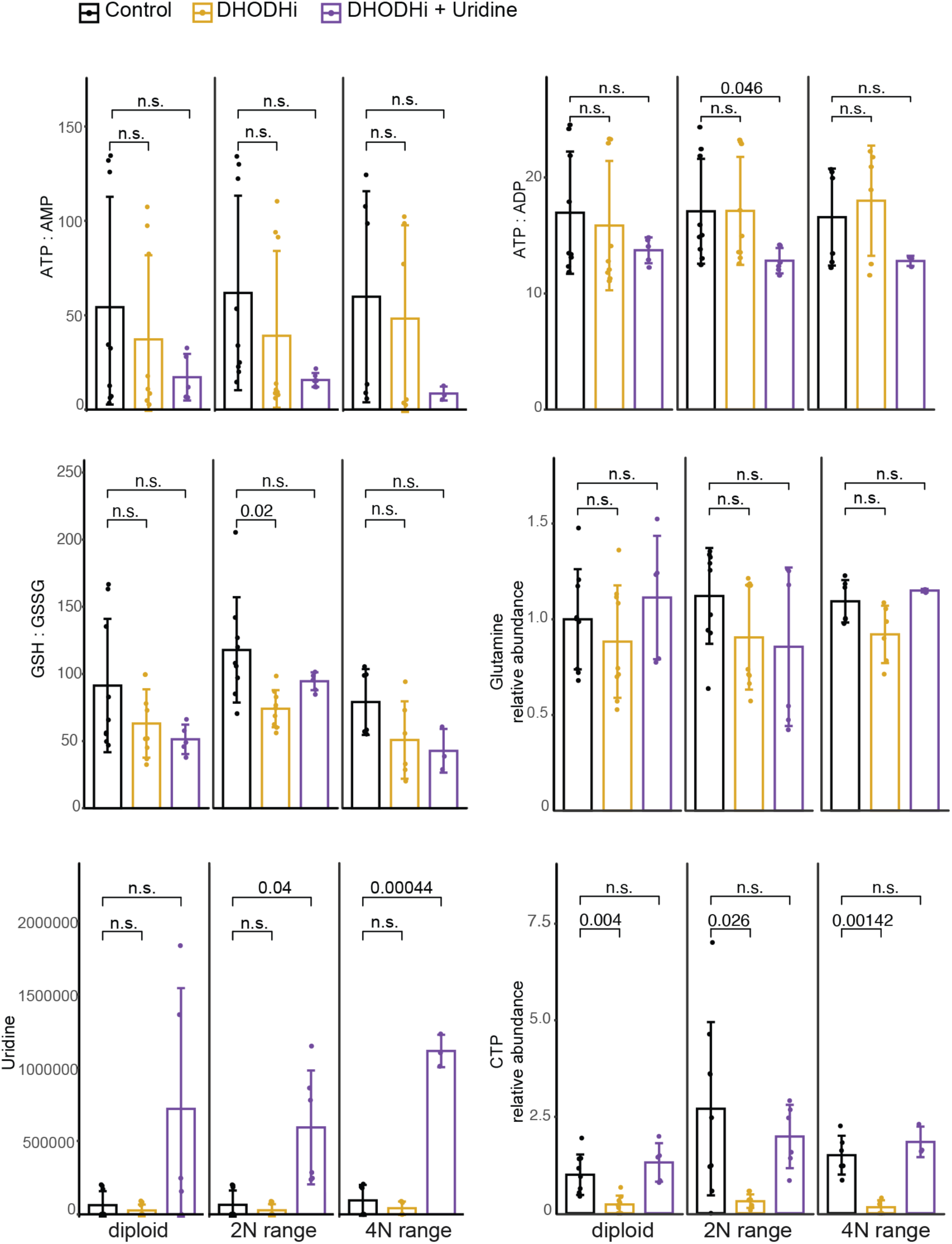
Metabolites beyond just pyrimidines are disrupted by DHODH inhibition in net-gain aneuploid HMECs. Bar graphs showing unchanged ratios of ATP::AMP or ATP::ADP metabolites and metabolite ratios in diploid, 2N-range aneuploid, and 4nN-range aneuploid cells under control conditions, treatment with DHODH inhibitor, and treatment with DHODH inhibitor and uridine rescue. P-values were calculated using T tests.

**SUPPLEMENT FIGURE 4:**
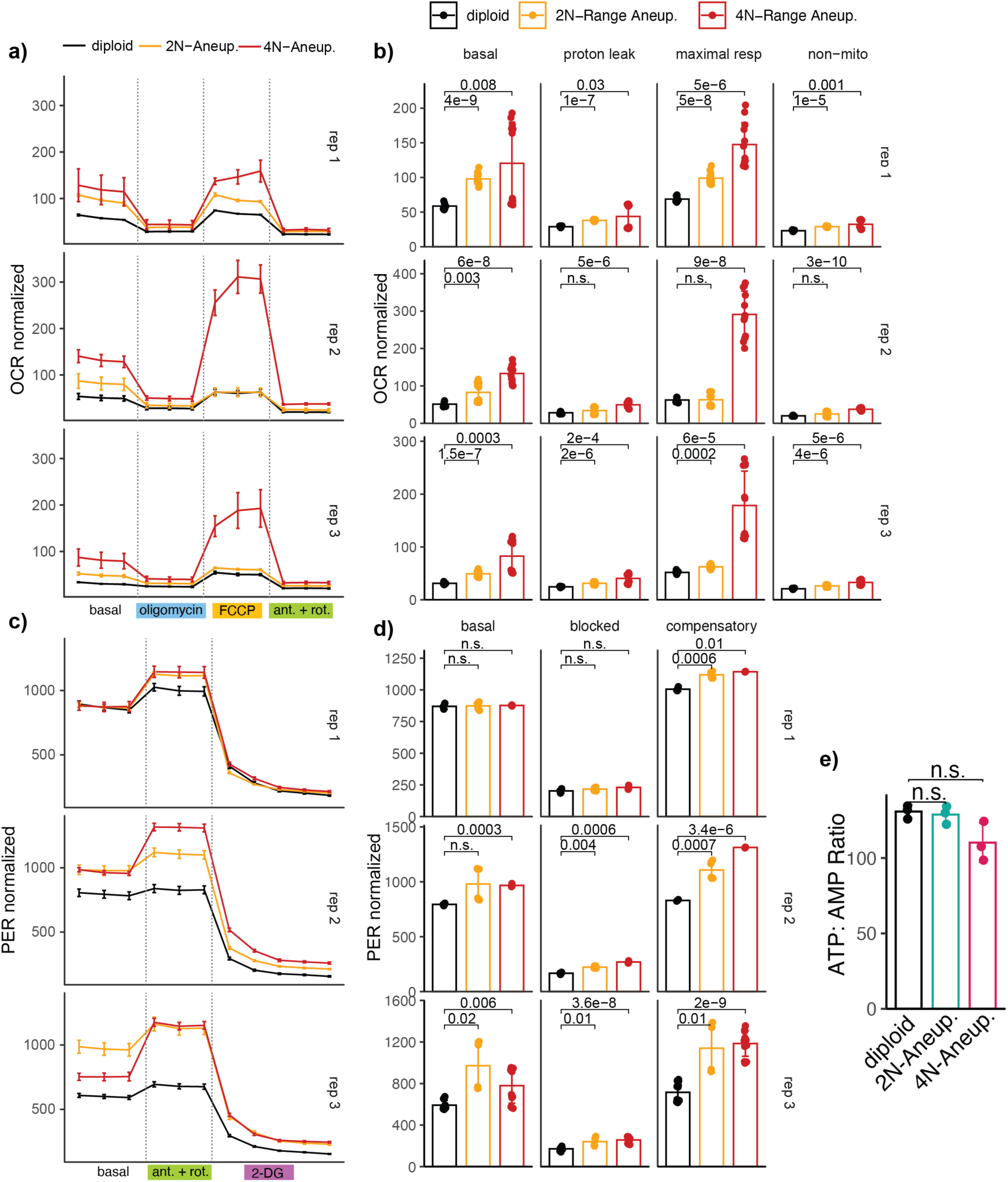
Individual replicate experiments of mitochondrial stress test and glycolytic rate assays. **a,** Oxygen consumption rates of three independent biological replicates of the Mitochondria Stress Test Assay using diploid, 2n-range aneuploid, and 4n-range aneuploid HMECs. Error bars indicate the standard deviation of technical replicate wells. **b,** Bar graphs showing the basal respiration, proton leak, maximal respiration, and non-mitochondrial respiration of diploid, 2n-range aneuploid, and 4n-range aneuploid HMECs across three replicates. P-values were calculated using T tests corrected for multiple hypothesis testing. **c,** Proton efflux rate of three independent biological replicates of the Glycolytic Rate Assay using diploid, 2n-range aneuploid, and 4n-range aneuploid HMECs. Error bars indicate the standard deviation of technical replicate wells. **d,** Bar graphs showing the basal, compensatory, and non-glycolytic proton efflux of diploid, 2n-range aneuploid, and 4n-range aneuploid HMECs across three replicates. P-values were calculated using T tests corrected for multiple hypothesis testing. **e,** Ratio of ATP:AMP in diploid, 2N-range, and 4N-range aneuploid HMECs under steady state conditions.

**SUPPLEMENT FIGURE 5:**
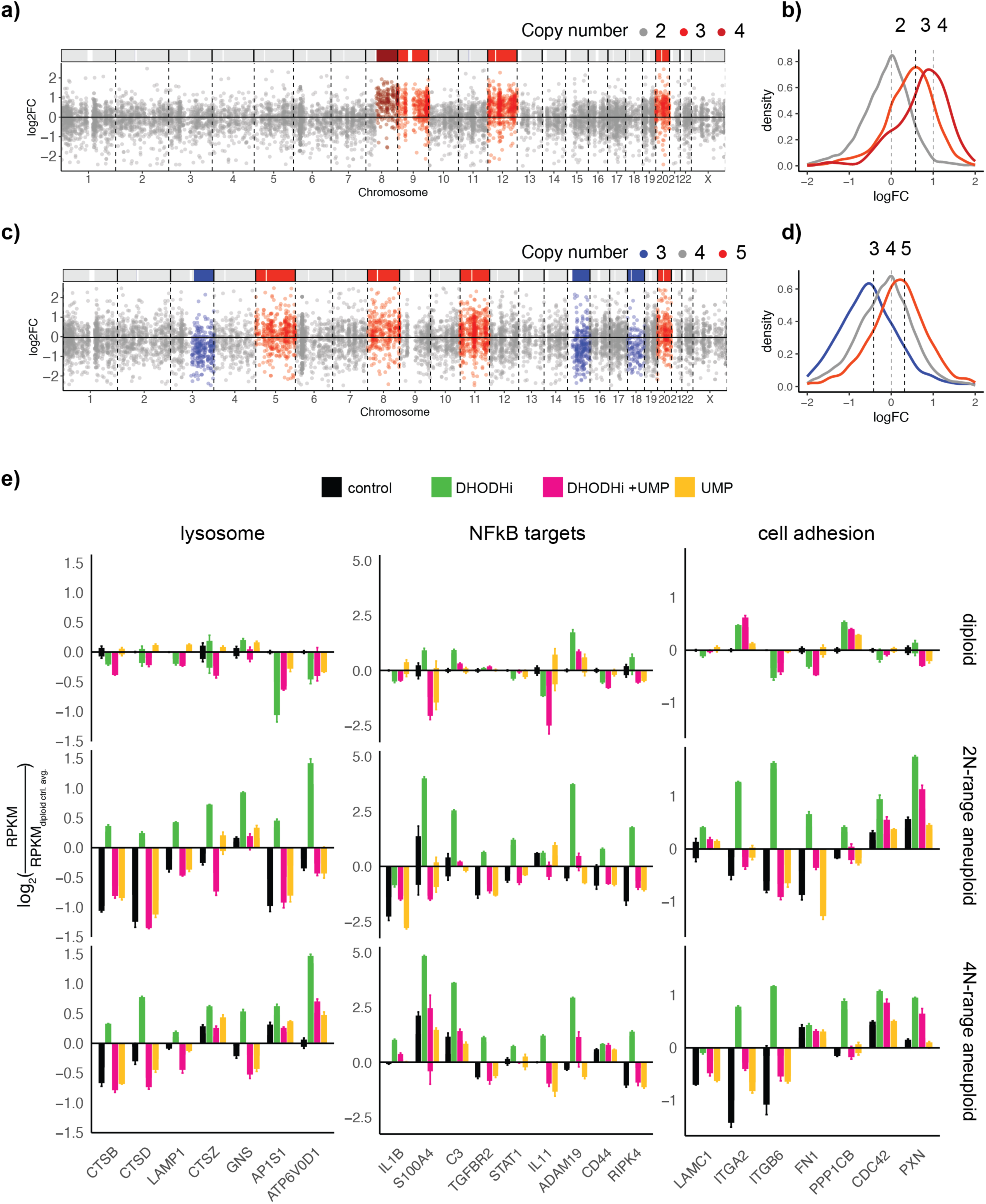
Aneuploidy-dependent gene expression changes. **a-d,** As a validation of the dataset shown in Fig. 5 a-b, we observe expected copy number-dependent gene expression changes when comparing aneuploid to diploid control RNA-seq data. Gene expression log_2_ fold change in a 2N-range aneuploid mutant relative to diploid controls plotted by genomic position **(a)** and as a density plot with respect to ploidy **(b)**. A similar genomic position plot **(c)** and density plot **(d)** for a 4N-range aneuploid HMEC line relative to diploid HMECs is shown. Copy number profiles for each line used are indicated by solid horizonal barplot across the top of each scatter plot. Genes are colored by gain/loss status. **e,** Bar graphs showing relative reads per kilobase per million reads of specific lysosome, NFkB, and cell adhesion genes in diploid, 2N-range aneuploid, and 4N-range aneuploid cells treated with the indicated conditions. These gene sets were upregulated in aneuploid cells but not diploid cells under DHODH inhibition.

**SUPPLEMENT FIGURE 6:**
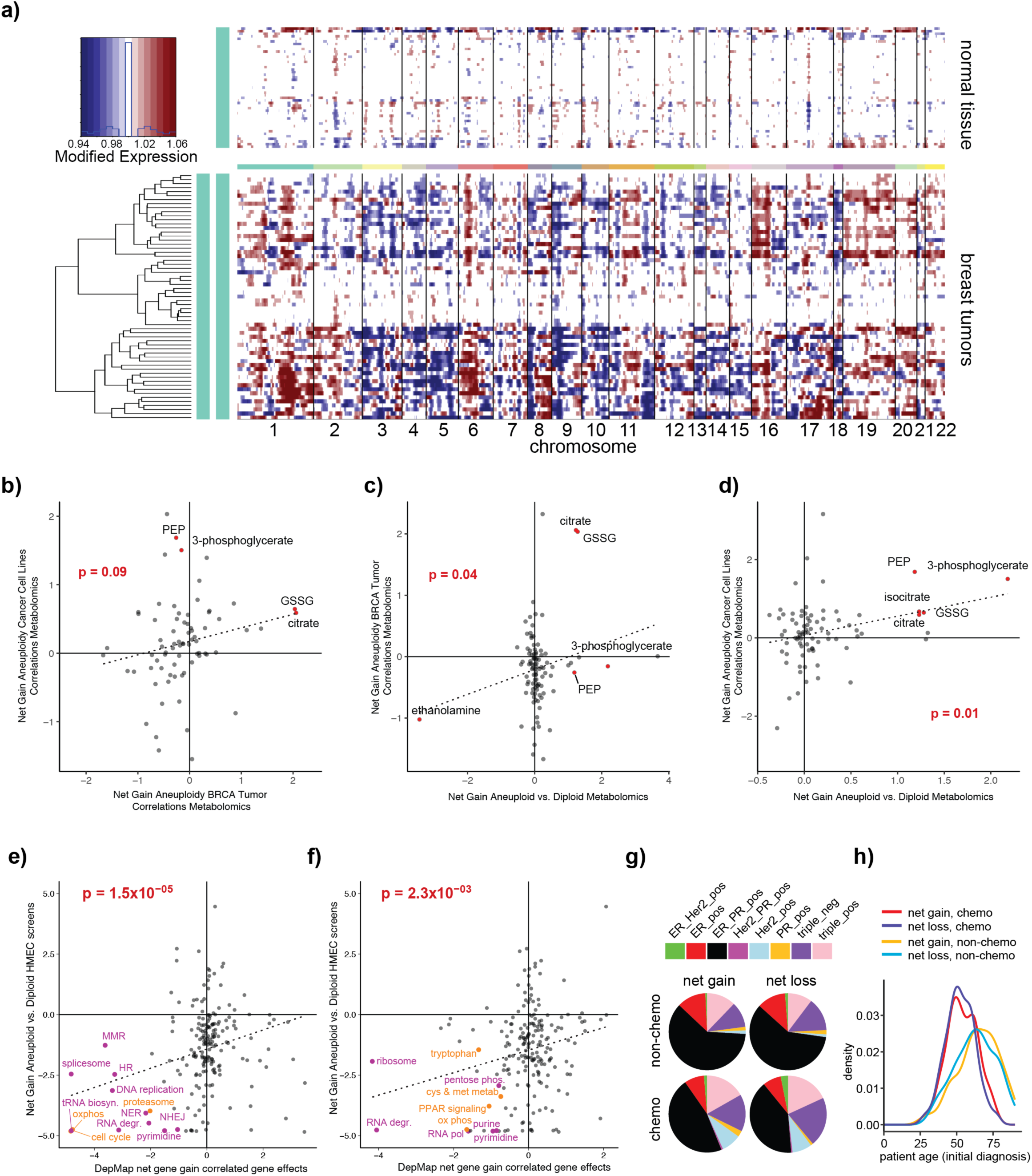
Net-gain aneuploidy is associated with metabolic phenotypes and is prognostic for response to DNA-damaging or anti-nucleotide chemotherapy in cancer. **a,** InferCNV^75^ output of breast cancer tumor sample RNA-seq dataset used to infer copy number status for metabolomic phenotype analysis. Samples used for the control group were derived from normal tissue. **b-d** Comparison between the metabolic profile associated with net-gain aneuploidy in this study (HMECs) and in a breast cancer metabolomics dataset (corresponding to the tumors in **(a)**) **(b)**, or in cancer cell line metabolomics datasets **(c)**, or between the tumor and cancer cell line datasets **(d)**. P-values are derived from linear regression analysis. Positive values indicate metabolites that are increased with net-gain aneuploidy; negative values indicate metabolites that are decreased with net-gain aneuploidy. **e, f,** Correlation between the pathway-level epistasis profile obtained from our HMEC net-gain aneuploidy CRISPR screens (y-axis) and the profile obtained from DepMap-based analysis of net-gain aneuploid pan-cancer cell lines (x-axis) **(e)** or just breast cancer cell lines (x-axis) **(f)**. Gene set enrichment analysis using KEGG gene sets of net-gain aneuploidy association rankings in both screens provided the scores for each pathway, which are calculated as the directional - log_10_(P-value) associated with pathway enrichment or depletion. Negative values indicate pathways that are synthetic-lethal with net-gain aneuploidy. The plotted P-value is derived from linear regression of the two profiles. **g,** Estrogen receptor (ER), Progesterone receptor (PR), and Her2 status (and combinations thereof) distributions among TCGA patients utilized for survival analysis in Fig. 6g show no significant difference between net gain and net aneuploidy loss tumor classification groups in the chemotherapy-treated (chemo) and non-chemotherapy-treated (non-chemo) cohorts. See Methods section for description of DNA-damaging/anti-nucleotide/anti-metabolite drug classes that form the “chemo” group in this study. **h,** Distributions of patient age at initial diagnosis within the groups/cohorts used in the survival analysis in Fig. 6g. There is no significant difference in age distribution between net gain and net aneuploidy loss tumor classification groups within each treatment cohort, though the chemo-treated cohort has a younger age distribution than the non-chemo cohort.

